# Association between Lifestyle Habits (Physical Exercise, Sleep, BMI, and Tobacco Consumption) and Academic Performance in Spanish University Students: A Cross-Sectional Study

**DOI:** 10.64898/2026.08.11.744105

**Authors:** Blanca Tardieu, Pablo Pons, Teresa López de Coca, Gonzalo Haro, Ana Benito, Pilar Sanfeliu

## Abstract

Healthy lifestyle habits are a key pillar of physical, psychological, and social well-being, yet university years are often marked by declines in sleep quality, physical activity, and other health behaviors that may affect academic performance. This cross-sectional study examined these associations in 390 Spanish university students, assessed via an online and in-person questionnaire covering academic trajectory (grade point average, credits passed, curricular progression), sleep quality (PSQI), physical activity (IPAQ), tobacco use, Body Mass Index (BMI), and sociodemographic data. Bivariate associations were tested with chi-square, Pearson correlation, t-tests, and ANOVA, followed by generalized linear models for significant variables. Sports participation, but not IPAQ-measured physical activity, was significantly associated with academic progression, defined as advancing one grade level per year. Sleep indicators showed mixed, partly counterintuitive associations: poorer nocturnal sleep was linked to better progression, whereas better sleep quality predicted a higher percentage of credits completed. BMI was a modest positive predictor of credit completion, and underweight students performed significantly worse than overweight or obese peers. Tobacco use showed no significant associations. These findings suggest that sleep-related behaviors, sports participation, and, to a lesser extent, BMI, are relevant correlates of academic progress, supporting university health policies addressing sleep, physical activity, and nutrition.

## 1. Introduction

Healthy lifestyle habits play a fundamental role in physical, mental, and social well-being. The World Health Organisation (WHO) defined a ‘Healthy Lifestyle’ (HL) as a way of living that reduces the likelihood of developing serious illnesses or premature death, offering more than just disease prevention (World Health Organisation, 1999). However, literature still debates which habits are the most decisive for such well-being. Nutrition, physical exercise, and tobacco abstinence have been highlighted by the WHO as dimensions that have a significant effect on the development of HL (World Health Organisation, 1999). Additionally, numerous studies have demonstrated correlations between sleep and various physical and mental health consequences, thus emphasising its importance in the development of HL (Jike et al., 2018).

During their university years, young adults undergo a transition that frequently entails negative changes in their lifestyle habits. Various studies have documented a decrease in physical activity, a decline in sleep quality, and an increase in the consumption of substances such as tobacco (Puente-Hidalgo et al., 2024). Research has also found that the worsening of these habits increases academic burnout and diminishes performance (Alexopoulou et al., 2024; Pastor et al., 2017).

The WHO identifies physical exercise as a fundamental pillar of a healthy lifestyle that contributes to general well-being by stimulating the body’s natural maintenance and recovery systems (World Health Organisation, 1999). It recommends engaging in moderate-intensity physical activity, defined as that which causes slight shortness of breath, for at least thirty minutes, five or more days a week. Physical exercise is another habit that often undergoes changes during the transition to university life: 42% of students reported exercising less frequently since starting university (Silliman et al., 2004). The most common reasons for this decline in physical exercise were a lack of time, motivation, and the mental energy to begin. A recent international systematic review found that, in general, it could be concluded that physical exercise has a moderate positive effect on academic performance (Rosales-Ricardo & Cáceres-Manzano, 2024). The review also highlighted that although some studies support this assertion, others fail to prove it. Within literature, specific inconsistencies lie in the lack of consensus on the types of physical activity associated with academic performance and the mediators behind this association. For example, some studies find that physical activity and strength training are both associated with academic performance (Reuter & Foster, 2021), whereas others only report modestly higher GPA for physical activity and not for strength training (Wald et al., 2014). Regarding the mediators, a recent study has found that physical exercise contributes to emotional regulation, attention and executive functions, key processes for academic functioning (Zhao et al., 2024). In the Spanish population in particular, negative associations have been found between academic stress and physical exercise. Regular physical activity helps reduce academic stress (Monserrat-Hernández et al., 2023), which might in turn lead to better academic performance.

Regarding sleep, a healthy sleep pattern can be approximated based on the criteria of the Pittsburgh Sleep Quality Index (PSQI) (Buysse et al., 1989): a nightly rest of more than 7 hours, where the majority of time in bed is spent sleeping, and it takes no longer than 30 minutes to fall asleep. It is also characterised by the absence of prolonged nighttime awakenings, use of sleep medication, sleep disorders, and negative impacts on daytime activities.

Lund et al. (2010) conducted a study with a sample of 1,125 students that found that over 60% had poor sleep quality according to the PSQI categorisation. The factors that most negatively impacted sleep were emotional and academic stress. This aligns with the findings of Puente-Hidalgo et al. (2024), who argued that the decline in healthy habits among university students increases as academic demands rise.. Furthermore, a study on Spanish university students found that during exam periods, students reported sleeping less than desired, and 61.3% of students believed that their performance would improve by getting more sleep (Suardiaz-Muro et al., 2023). The study also found a positive association between sleep quality and academic performance. A systematic review on the relationship between sleep and academic performance suggests that daytime sleepiness and drowsiness worsen academic performance (Suardiaz-Muro et al., 2020). Research has also established that sleep disturbances are associated with cognitive decline and, in university students in particular, are negatively associated with different cognitive functions (language, visual-perceptual ability, verbal and visuospatial memory and attention and concentration), which could be behind the association with academic performance (Xu et al., 2025).

Regarding nutrition and nutritional status, the World Health Organisation (WHO) defines obesity as a condition characterised by excessive body fat that compromises health and well-being. For its quantification, the WHO recommends the use of the Body Mass Index (BMI), calculated as the ratio of weight (kg) to height squared (m²) (Lemos et al., 2001). However, although BMI is a widely utilised tool in epidemiological research, it possesses inherent limitations in accurately reflecting body composition and fat distribution, factors that are critical for understanding its functional and cognitive implications. (Wells & Fewtrell, 2006; Romero-Corral et al., 2008; Rothman, 2008)

In this context, university students represent a particularly vulnerable demographic.They are in a transitional life stage characterised by the adoption of habits that often include unhealthy dietary patterns and sedentary lifestyles. These behaviours have a potential impact on both health and academic performance (Ramírez et al., 2019). The persistence of such habits over time may consolidate into risk factors for the development of chronic diseases (Sánchez-Ojeda & Luna-Bertos, 2015). Furthermore, evidence suggests that obesity among university students is associated with increased susceptibility to psychological problems, such as anxiety, depression, and greater social isolation, variables that can negatively mediate academic achievement (Wehigaldeniya et al., 2017).

Concurrently, numerous studies in adult populations have linked obesity to poorer cognitive functioning, including deficits in visual and verbal memory, attention, decision-making, and inhibitory control, as well as neurostructural alterations in key regions such as the orbitofrontal cortex (Boeka & Lokken., 2008; Weller et al., 2008; Reinert et al., 2013; Prickett et al., 2015). Nevertheless, the extrapolation of these findings to the university population remains limited, highlighting the need to further investigate the impact of obesity on psychological and cognitive functioning during this developmental stage. Furthermore, other studies have explored variables that could address nutrition more directly in Spain, such as adherence to the Mediterranean diet (e.g., López-Gil et al., 2023; Alfaro-González et al., 2024; Monserrat-Hernández et al., 2023). However, studies on the link between academic performance and BMI are scarcer in this population.

The WHO identifies smoking as the greatest self-imposed health risk (World Health Organisation, 1999). It contributes to the development of respiratory diseases, coronary heart disease, and various types of cancer. The WHO emphasises the importance of quitting smoking at any age to significantly reduce health risks. In addition to these health risks, it is relevant to examine whether tobacco consumption might also influence students’ academic performance. This habit is particularly significant in this population, as 82.6% of smokers begin smoking between the ages of 15 and 24, and students from various universities identify university campuses as the second most common space where they consume tobacco products and where they are most exposed to secondhand smoke Teixeira-da-Costa et al., (2022). A recent study in Spanish university students found that those who have failing grade averages consumed more tobacco than those with outstanding grade averages (Llorent-Bedmar et al., 2023). Furthermore, another recent study in Spanish high school students found that the positive association between the Mediterranean diet (considered a healthy diet and associated with better nutrition) and academic performance could be moderated by the use of tobacco (López-Gil et al., 2023). Additionally, tobacco consumption is often associated with other risky behaviours such as alcohol consumption and cannabis use, which also collectively contribute to poorer academic outcomes (Llorent-Bedmar et al., 2023; Páramo et al., 2020). Other possible factors that mediate this relationship have been explored in the Spanish population, such as emotional intelligence or social support; they were not found to be significant (Rodríguez-Sáez et al., 2021). Thus, this study aims to provide supporting evidence for this association between tobacco use and academic performance.

Factors such as sleep, physical activity, dietary habits, and tobacco avoidance, constitute the foundation of a healthy lifestyle. All four undergo a significant transition and often a decline during the university years and have been identified as influential factors in academic or cognitive performance. Various studies have investigated multiple health habits simultaneously to determine the significance of lifestyle choices in enhancing academic performance; for instance, Maniaci et al. (2023) examined these associations among Italian adolescents.

This study simultaneously examines four lifestyle habits in Spanish university students to explore their interactions and relationship with academic performance. By doing so, it seeks to reconcile gaps within the literature and provide universities with actionable insights for developing programs that promote healthier lifestyles among students. The hypotheses are: 1) the amount of physical exercise will be positively associated with academic performance; 2) healthy sleep habits will be positively associated with academic performance; 3) there will be a significant relationship between BMI and academic performance; and 4) tobacco consumption will be associated with academic performance.

## 2. Materials and Methods

### 2.1. Study Design and Participants

The study is observational and cross-sectional, with descriptive and analytical components. The sample size was determined using G*Power software (version 3.1.9.4). Results showed that a sample of 382 is required to compare two independent groups using a t-test with an effect size of 0.3, alpha of 0.05, and power of 0.90. Additionally, a sample of 92 is required to calculate correlations, and 38 is required to perform multiple linear regression with these same parameters.

The inclusion criteria were a) being a university student in their 2nd year or higher; b) being between 18 and 25 years of age; and c) signing the informed consent form. Students in the first academic year were excluded because they had not yet completed a full university assessment cycle at the time data collection took place (between September and December 2024 before the first end-of-semester examination period in Spain).. This criterion was used to contextualize academic results based on participants’ prior experience with university-level academic and examination standards. The age range was selected to focus on young adults in the typical university stage and to reduce heterogeneity related to more diverse academic and life-course trajectories. The exclusion criteria were a) students from Cardenal Herrera-CEU University (Valencia, Spain), to avoid the ethical conflict associated with using a captive sample; and b) exchange students, including both international students and those who had completed a study abroad program of at least one academic year.

In the 2023–2024 academic year, the University of Valencia (UV) had approximately 65,800 students, while the Polytechnic University of Valencia (UPV) had around 31,200 students. Based on these population figures, the sample selection was conducted using a proportional criterion, to respect the relative distribution of students from both institutions. Thus, approximately 63% of the sample corresponded to UV students and 33.8% to UPV students, to reflect more accurately the real composition of the reference university population.

Data collection took place between September and December 2024. In the first phase, data was collected in person in public areas of Valencia located near different universities. Recruitment was conducted at street level, approaching potential participants in the vicinity of university campuses. Questionnaires were administered to those who agreed to participate and met the eligibility criteria. The first phase followed a street-intercept approach among students present in these areas. Subsequently, a second data collection phasewas conducted virtually to optimise the process. A link to the questionnaire was distributed among university students to encourage greater participation and broaden the reach of the study. To ensure data quality, both recruitment procedures used the same questionnaire and eligibility criteria.

### 2.2. Instruments

- **Sleep Quality:** The Pittsburgh Sleep Quality Index (PSQI) (Buysse et al., 1989) was used in its Spanish adaptation (Royuela Rico & Macías Fernández, 1997). It assesses sleep quality over the past month, providing a global score (higher scores indicate poorer sleep quality) and seven specific components. For descriptive and comparative analysis, participants were grouped according to the conventional PSQI cut-off: ≤ 5 points, good sleep quality; > 5 points, poor sleep quality. Additionally, the Jenkins Sleep Scale-4 (JSS-4) (Jenkins et al., 1988), validated in Spanish by Villarreal-Zegarra et al. (2022), was administered. Scores >12 indicate poor sleep quality.
- **Physical Activity:** The International Physical Activity Questionnaire (IPAQ) (Craig et al., 2003) was administered in its Spanish version (Roman-Viñas et al., 2010). It evaluates the frequency and duration of light, moderate, and vigorous physical activities, including information on participation in sports and other regular physical activities, which was used to identify the type and intensity of activity performed by each participant and to calculate energy expenditure in METs (Metabolic Equivalent of Task). Responses were scored following the standard protocol of the IPAQ Research Commitee (2005), which classifies participants into three categories (low, moderate, or high) based on frequency, duration, and total MET-minutes/week. Specifically, the “high” category includes participants performing vigorous activity on at least 3 days, achieving a minimum of 1500 MET-min/week, or 7 or more days of any combination of walking, moderate, or vigorous activity achieving at least 3000 MET-min/week.
- **Tobacco Consumption:** Two ad hoc items were included: (a) presence or absence of tobacco use, and (b) number of cigarettes smoked per day.
- **Complementary Variables:** Sociodemographic data (age, sex, field of study, academic year, parental education level, type of residence) and physical health data (body mass index, presence of medical pathology) were collected.
- **Academic Performance:** Three indicators were considered: (a) grade point average (GPA) of the degree, (b) percentage of credits passed, and (c) academic progress (whether the student is progressing normally, defined as not having repeated subjects or academic years).

### 2.3. Data Analysis and Ethical Considerations

Data analysis was conducted using IBM SPSS Statistics (v.29). Following an exploratory and descriptive analysis of the variables, the relationships between the variables of interest were examined. The central limit theorem allows us to assume normality in samples with n ≥ 30 (Kwak & Kim, 2017), so parametric tests were used. Specifically, a chi-square test was employed for categorical variables, while Pearson’s correlation coefficient (r) was used for quantitative variables. To compare quantitative variables between groups, an independent samples t-test was applied for two-group comparisons, and an analysis of variance (ANOVA) was utilised for variables with three or more categories.

Subsequently, variables demonstrating significant associations with each academic performance indicator in the bivariate analyses were incorporated into generalised linear models (GLM) to evaluate their predictive capacity. Each predictor was first examined in an unadjusted model. Adjusted models were then constructed separately for each outcome, including all variables that showed statistically significant associations with that specific outcome. In addition, potential interaction effects between key independent variables (e.g., gender, physical activity, and sleep indicators) and sociodemographic characteristics were explored within the GLM framework. Interaction terms were included to assess whether the associations between lifestyle factors and academic performance varied across subgroups. No statistically significant interaction effects were identified, suggesting that the observed relationships were relatively consistent across the groups analysed. Since the GLM procedure does not provide the beta values, the variables were standardized to calculate them, although they must be interpreted with caution due to the presence of categorical variables.

For academic progress (on track), the adjusted model included father’s education level, sports participation, nocturnal sleep hours, daytime sleep hours, PSQI score, and Jenkins sleep category. For the percentage of credits completed, the model included PSQI score and BMI. For GPA, the model included age, gender, field of study, and academic year. The “academic year” variable was excluded from the predictive model for the “year-by-year” indicator to prevent collinearity issues.

Finally, SmartPLS 4 (Ringle, Wende, & Becker, 2024) was used to estimate a path model examining the relationships among the study variables. Associations were evaluated using standardized β coefficients and coefficients of determination (R²). Statistical significance was assessed using the consistent PLS-SEM bootstrapping procedure with 5,000 bootstrap subsamples.

The study was conducted in accordance with local ethical guidelines and European regulations regarding the protection of research participants, complying with the principles of the Declaration of Helsinki and the recommendations of the Ministry of Health for clinical studies. The protocol was approved by the Ethics and Biomedical Research Committee of Universidad Cardenal Herrera-CEU (Ref: CEEI23/440, 19 February 2024).

During the preparation of this manuscript, the authors used a generative AI tool (ChatGPT) to assist with language polishing and translation. The authors have reviewed and edited the output and take full responsibility for the content of this publication.

## 3. Results

### 3.1. Sample characteristics

Table 1 provides a detailed description of the study sample.

**Table 1.**
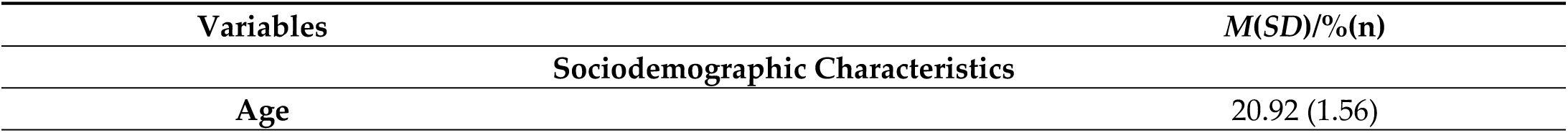
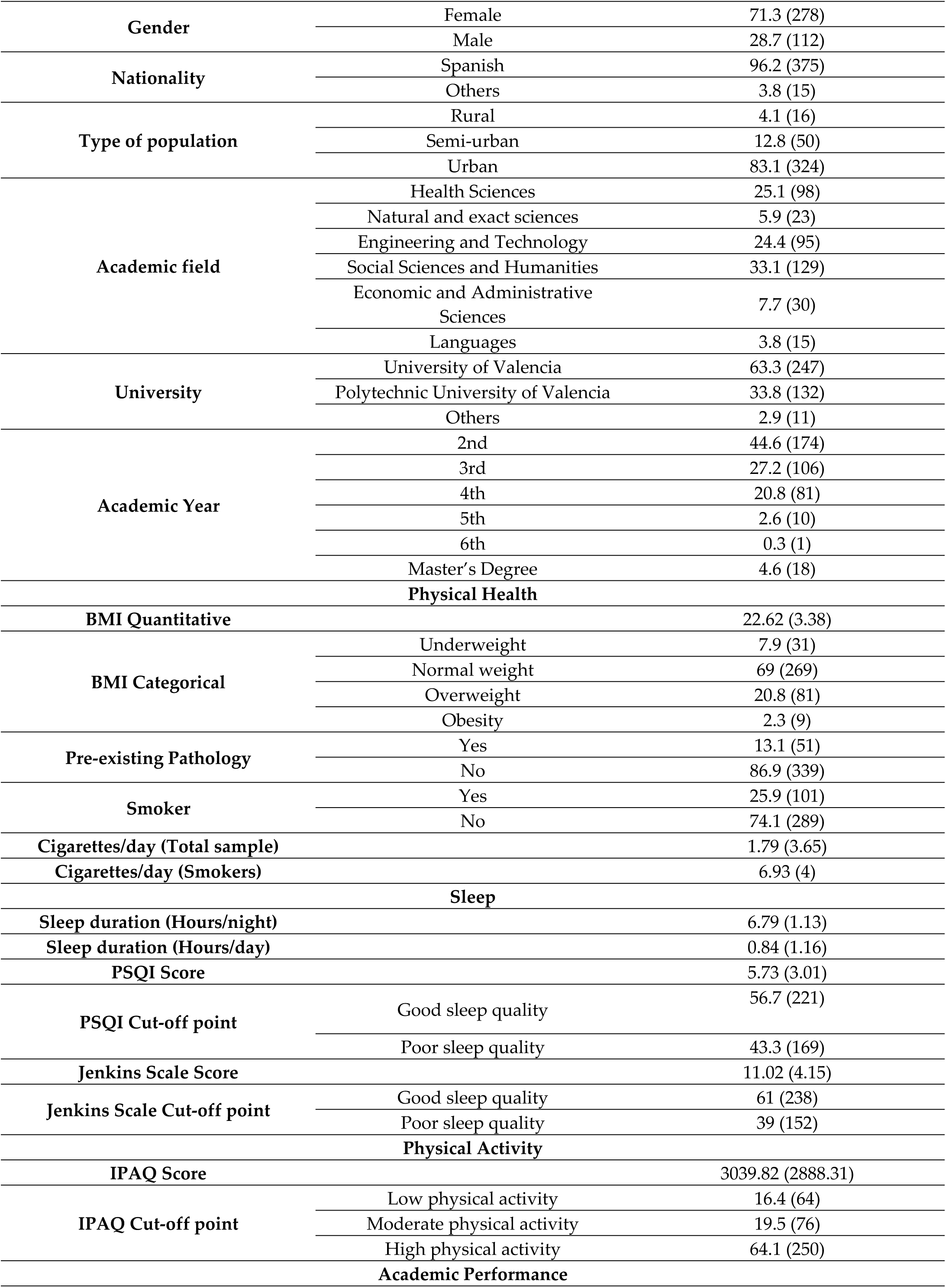
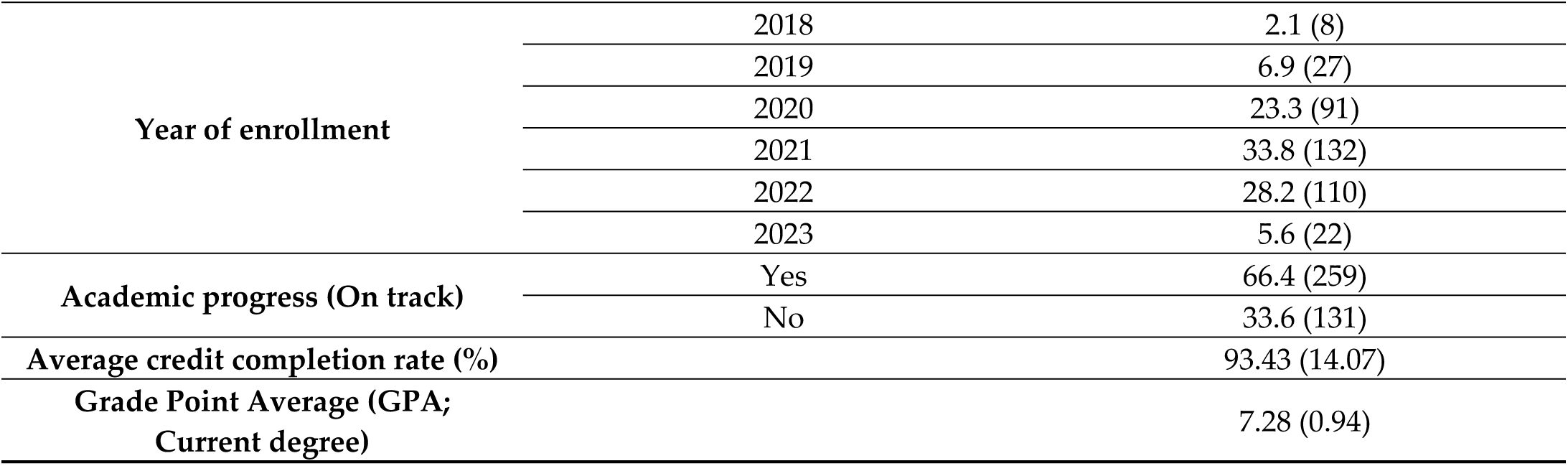
Descriptive statistics of the sample.

| Variables | *M(SD)/%(n)* |
| --- | --- |
| **Sociodemographic Characteristics** |  |
| Age | 20.92 (1.56) |

|  |  |  |
| --- | --- | --- |
| **Gender** | Female | 71.3 (278) |
|  | Male | 28.7 (112) |
| **Nationality** | Spanish | 96.2 (375) |
|  | Others | 3.8 (15) |
| **Type of population** | Rural | 4.1 (16) |
|  | Semi-urban | 12.8 (50) |
|  | Urban | 83.1 (324) |
| **Academic field** | Health Sciences | 25.1 (98) |
|  | Natural and exact sciences | 5.9 (23) |
|  | Engineering and Technology | 24.4 (95) |
|  | Social Sciences and Humanities | 33.1 (129) |
|  | Economic and Administrative Sciences | 7.7 (30) |
|  | Languages | 3.8 (15) |
| **University** | University of Valencia | 63.3 (247) |
|  | Polytechnic University of Valencia | 33.8 (132) |
|  | Others | 2.9 (11) |
| **Academic Year** | 2nd | 44.6 (174) |
|  | 3rd | 27.2 (106) |
|  | 4th | 20.8 (81) |
|  | 5th | 2.6 (10) |
|  | 6th | 0.3 (1) |
|  | Master's Degree | 4.6 (18) |
| **Physical Health** |  |  |
| **BMI Quantitative** |  | 22.62 (3.38) |
| **BMI Categorical** | Underweight | 7.9 (31) |
|  | Normal weight | 69 (269) |
|  | Overweight | 20.8 (81) |
|  | Obesity | 2.3 (9) |
| **Pre-existing Pathology** | Yes | 13.1 (51) |
|  | No | 86.9 (339) |
| **Smoker** | Yes | 25.9 (101) |
|  | No | 74.1 (289) |
| **Cigarettes/day (Total sample)** |  | 1.79 (3.65) |
| **Cigarettes/day (Smokers)** |  | 6.93 (4) |
| **Sleep** |  |  |
| **Sleep duration (Hours/night)** |  | 6.79 (1.13) |
| **Sleep duration (Hours/day)** |  | 0.84 (1.16) |
| **PSQI Score** |  | 5.73 (3.01) |
| **PSQI Cut-off point** | Good sleep quality | 56.7 (221) |
|  | Poor sleep quality | 43.3 (169) |
| **Jenkins Scale Score** |  | 11.02 (4.15) |
| **Jenkins Scale Cut-off point** | Good sleep quality | 61 (238) |
|  | Poor sleep quality | 39 (152) |
| **Physical Activity** |  |  |
| **IPAQ Score** |  | 3039.82 (2888.31) |
| **IPAQ Cut-off point** | Low physical activity | 16.4 (64) |
|  | Moderate physical activity | 19.5 (76) |
|  | High physical activity | 64.1 (250) |
| **Academic Performance** |  |  |
| Year of enrollment | 2018 | 2.1 (8) |
|  | 2019 | 6.9 (27) |
|  | 2020 | 23.3 (91) |
|  | 2021 | 33.8 (132) |
|  | 2022 | 28.2 (110) |
|  | 2023 | 5.6 (22) |
| Academic progress (On track) | Yes | 66.4 (259) |
|  | No | 33.6 (131) |
| Average credit completion rate (%) |  | 93.43 (14.07) |
| Grade Point Average (GPA; Current degree) |  | 7.28 (0.94) |

The sample comprised 390 university students (71.3% female; *M* = 20.9 years, *SD* = 1.56). The participants were predominantly of Spanish nationality (96.2%) and resided in urban areas (83.1%). Regarding academic disciplines, the most represented fields were Social Sciences and Humanities (33.1%), followed by Health Sciences (25.1%).

In terms of health status, the body mass index (BMI) was *M* = 22.6 (*SD* = 3.38), with 69% of the participants falling within the normal weight range. Additionally, 25.9% of the sample identified as smokers (*M* = 6.9 cigarettes per day among smokers). Regarding physical activity, 64.1% demonstrated a high level of physical activity according to the IPAQ, and 64.9% reported engaging in regular sports practice.

Concerning sleep quality, 43.3% of participants exhibited poor sleep quality as measured by the PSQI and 39% according to the Jenkins Sleep Scale.

Finally, overall academic performance was high, with a mean grade of *M* = 7.28 out of 10 (*SD* = 0.94) and an average credit completion rate of 93.43%. Furthermore, 66.4% of the sample was up to date with their curriculum (completing one academic year per calendar year).

### 3.2. Sociodemographic Variables and Academic Performance

Given the large sample size, effect sizes were systematically interpreted alongside p-values to assess the practical significance of the findings. Overall, most effects ranged from small to moderate according to conventional benchmarks (Cohen, 1988).

Table 2 presents the association between sociodemographic variables and academic performance.

**Table 2.**
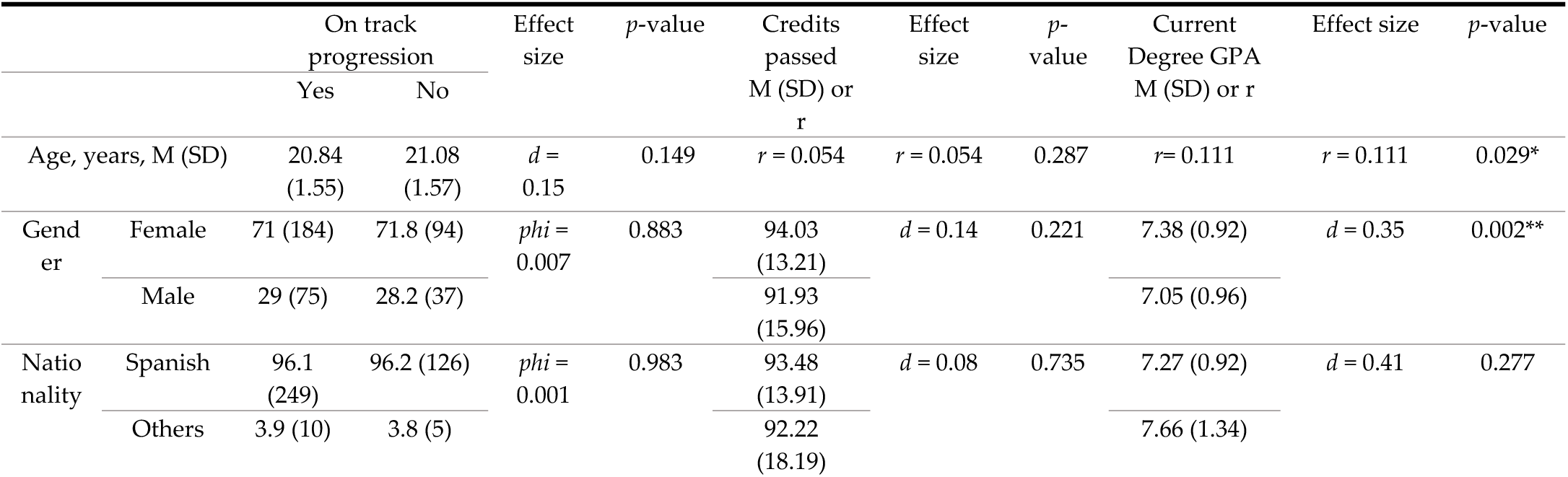
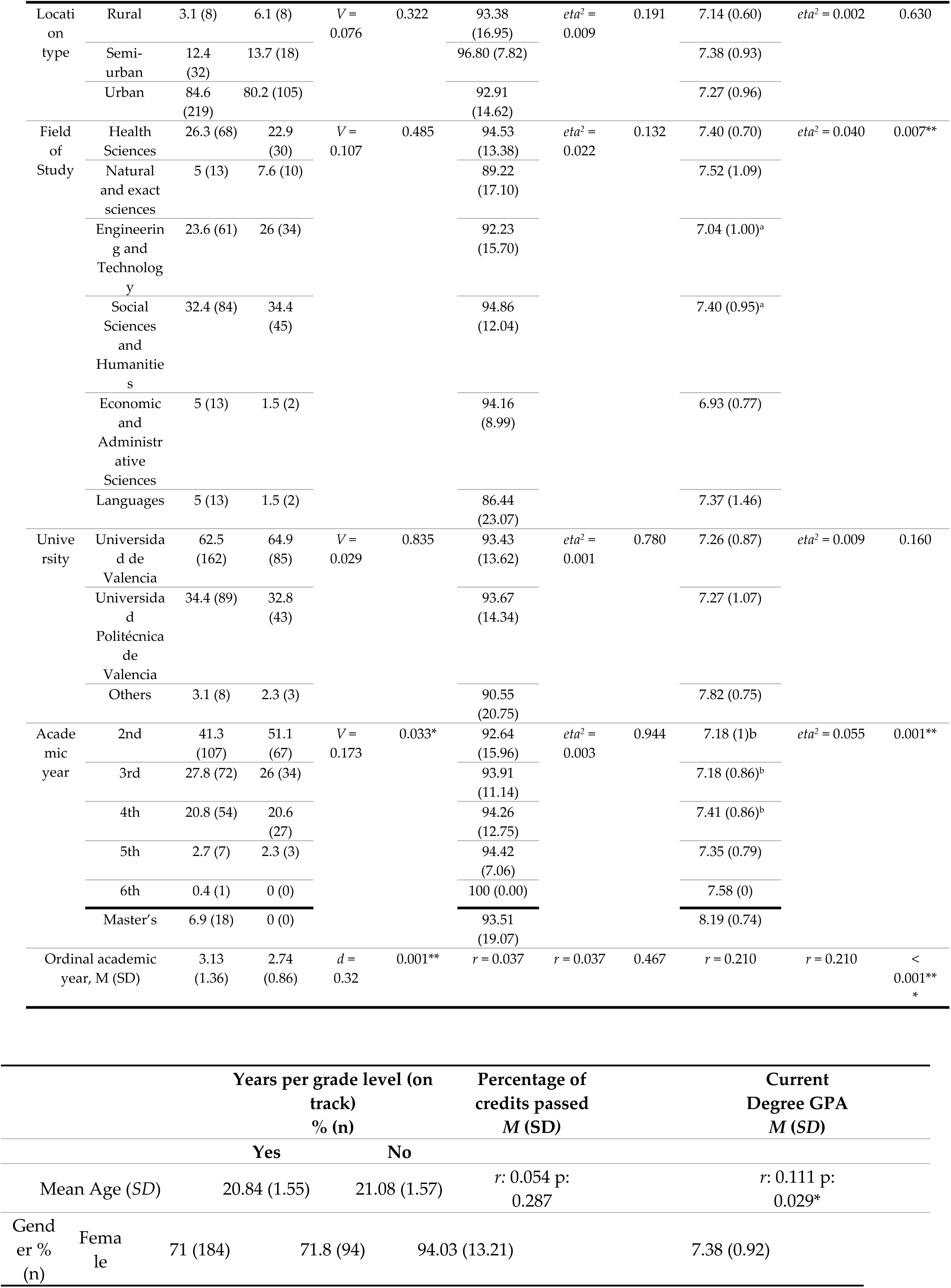
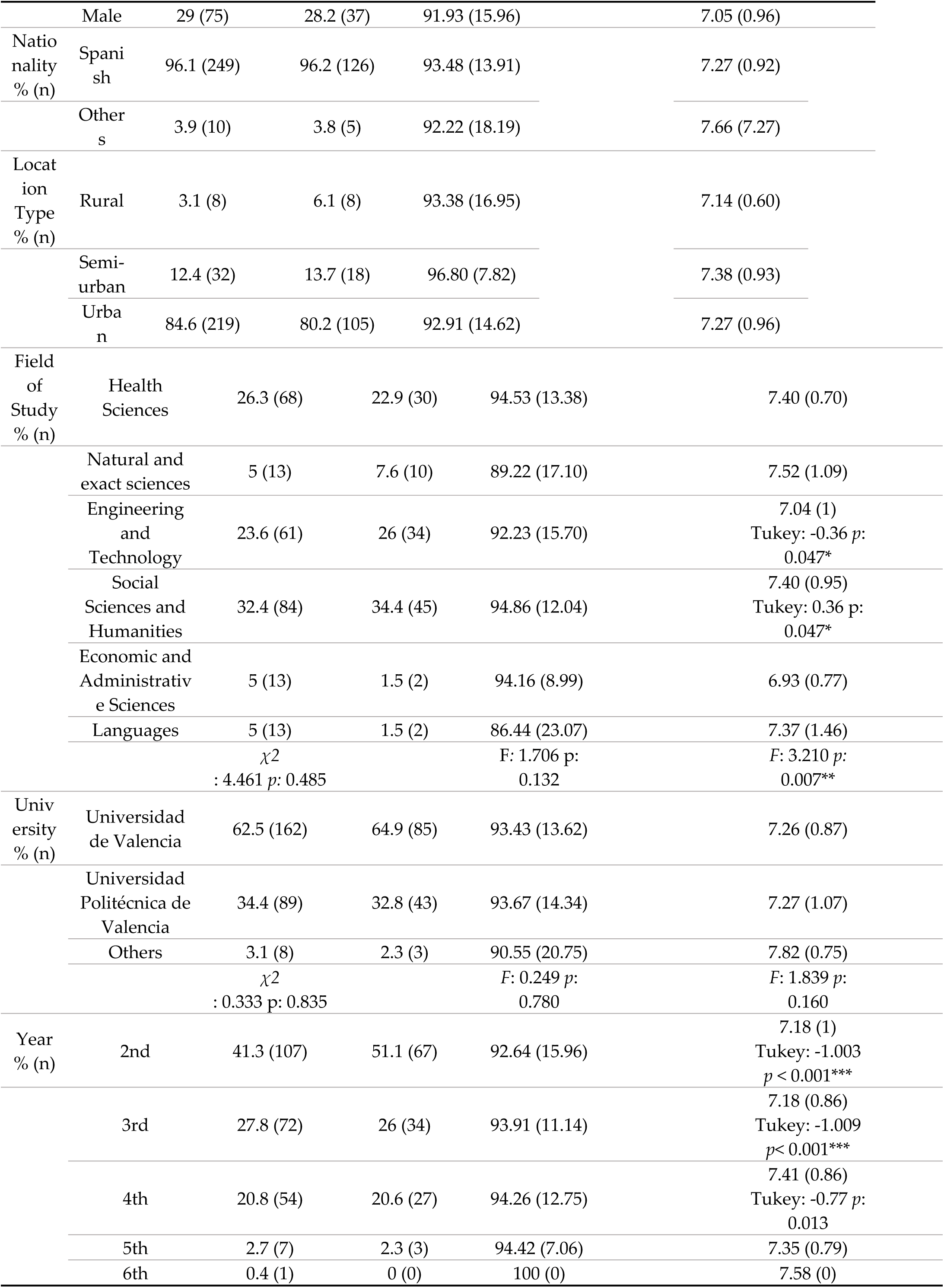
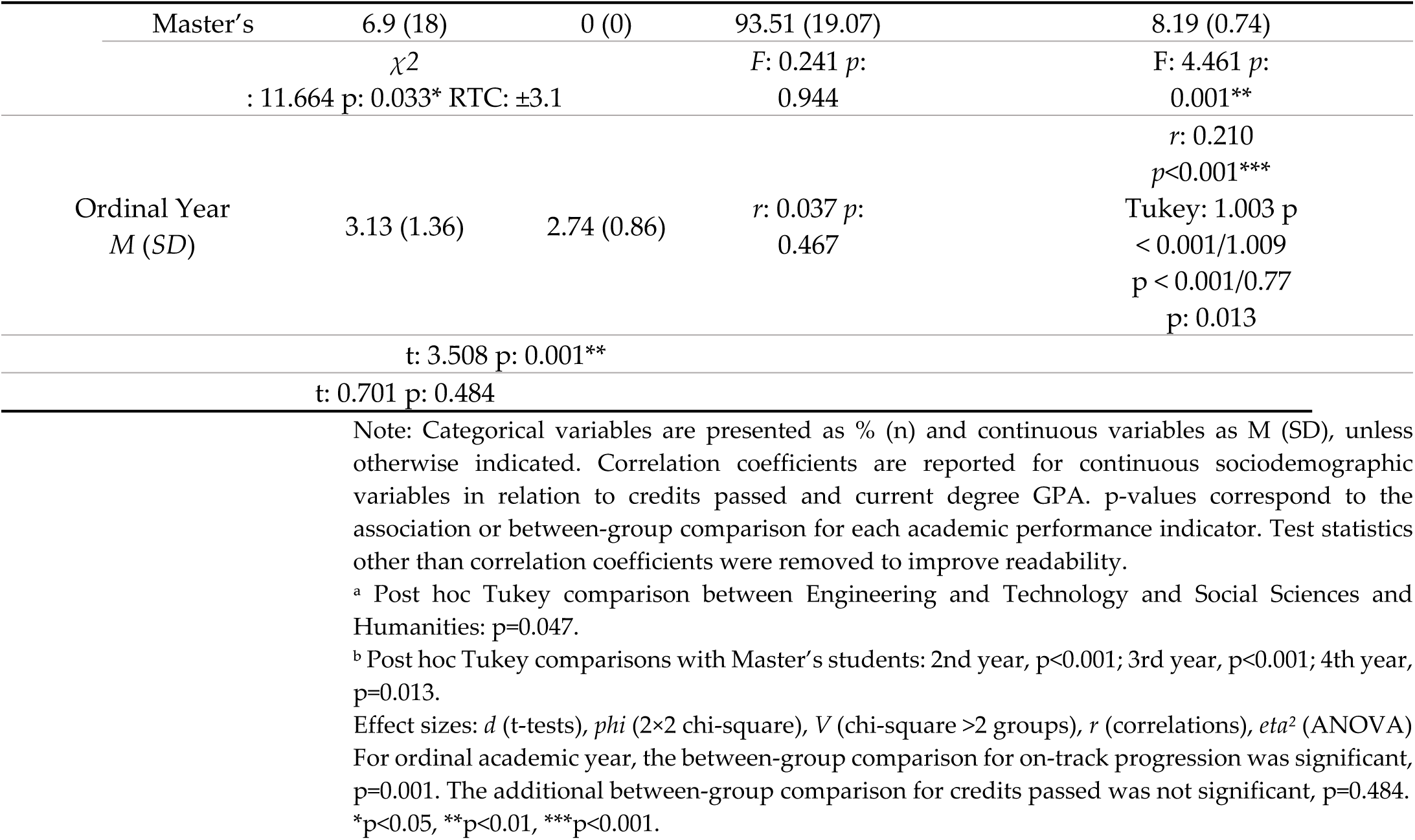
Relationship between sociodemographic variables and academic performance.

A weak positive correlation was observed between age and GPA (*r* = .111, *p* = .029), suggesting that older students tended to demonstrate slightly higher performance. Furthermore, significant gender differences were observed in GPA (*t* = 3.16, *p* = .002), with women achieving higher GPA than men. Notably, the effect size was small to moderate (*d* = 0.35), indicating that this difference is not only statistically significant but also meaningful.

The field of study showed a statistically significant association with GPA (F = 3.210, *p* = .007). However, the effect size was small (*eta*² = 0.040), suggesting limited practical relevance despite statistical significance. Post hoc comparisons indicated that students in Social Sciences and Humanities scored higher than those in Engineering and Technology. (*p* = .047).

The academic year correlated positively with both GPA (*r* = .210, *p* < .001), reflecting a small-to-moderate effect size, and academic progress (*t* = 3.51, *p* = .001), also with a small-to-moderate effect size (*d* = 0.32).

Regarding academic progress (years per grade level), an association was found with the current year of study (*χ2*= 11.664, *p* = .033), with a small effect size (*V* = 0.173). Students in higher-level years exhibited significantly higher GPAs, with notable differences between the second and fourth years (*p* = .013) and at the postgraduate level (*p* < .001). No significant associations were found for nationality, population type, or university of origin.

### 3.3. Physical Activity and Academic Performance

Participation in sports was significantly associated with academic progress (*χ*² = 7.24, *p* = .007), with a small effect size (*phi* = 0.136). Students engaged in sports demonstrated a higher probability of progressing through their curriculum without repeating a year (71.1%) compared to those who did not participate (57.7%), with adjusted standardized residuals (ASR = ±2.7).

Conversely, no significant associations were observed between physical activity levels as measured by the IPAQ (total score or categorical levels) and mean grade point average or the percentage of credits completed (*p* > .05).

Detailed results are available in Table 3.

**Table 3.**
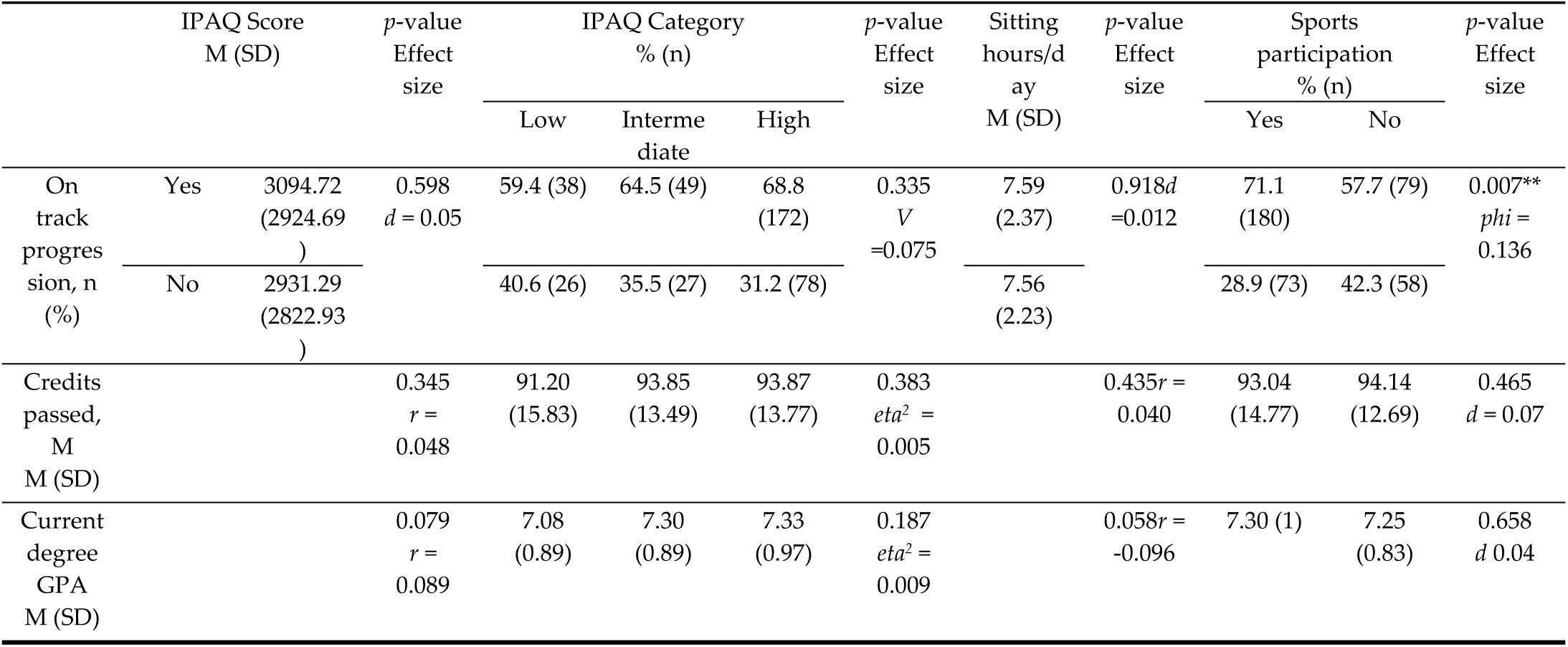
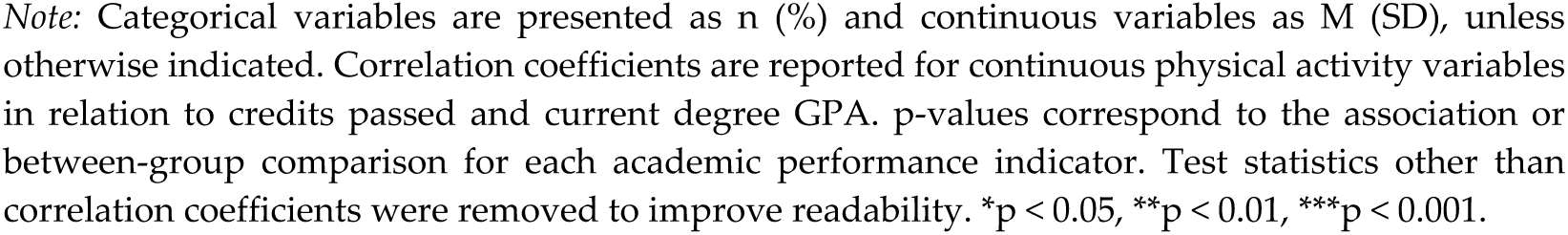
Relationship between physical activity and academic performance.

|  |  | IPAQ Score<br>M (SD) | *p*-value<br>Effect<br>size | IPAQ Category<br>% (n) |  |  | *p*-value<br>Effect<br>size | Sitting<br>hours/d<br>ay<br>M (SD) | *p*-value<br>Effect<br>size | Sports<br>participation<br>% (n) |  | *p*-value<br>Effect<br>size |
| --- | --- | --- | --- | --- | --- | --- | --- | --- | --- | --- | --- | --- |
|  |  |  |  | Low | Interme<br>diate | High |  |  |  | Yes | No |  |
| On<br>track<br>progres<br>sion, n<br>(%) | Yes | 3094.72<br>(2924.69<br>) | 0.598<br>*d* = 0.05 | 59.4 (38) | 64.5 (49) | 68.8<br>(172) | 0.335<br>*V*<br>=0.075 | 7.59<br>(2.37) | 0.918*d*<br>=0.012 | 71.1<br>(180) | 57.7 (79) | 0.007**<br>*phi* =<br>0.136 |
|  | No | 2931.29<br>(2822.93<br>) |  | 40.6 (26) | 35.5 (27) | 31.2 (78) |  | 7.56<br>(2.23) |  | 28.9 (73) | 42.3 (58) |  |
| Credits<br>passed,<br>M<br>M (SD) |  |  | 0.345<br>*r* =<br>0.048 | 91.20<br>(15.83) | 93.85<br>(13.49) | 93.87<br>(13.77) | 0.383<br>*eta*<sup>2</sup> =<br>0.005 |  | 0.435*r* =<br>0.040 | 93.04<br>(14.77) | 94.14<br>(12.69) | 0.465<br>*d* = 0.07 |
| Current<br>degree<br>GPA<br>M (SD) |  |  | 0.079<br>*r* =<br>0.089 | 7.08<br>(0.89) | 7.30<br>(0.89) | 7.33<br>(0.97) | 0.187<br>*eta*<sup>2</sup> =<br>0.009 |  | 0.058*r* =<br>-0.096 | 7.30 (1) | 7.25<br>(0.83) | 0.658<br>*d* 0.04 |

Note: Categorical variables are presented as n (%) and continuous variables as M (SD), unless otherwise indicated. Correlation coefficients are reported for continuous physical activity variables in relation to credits passed and current degree GPA. p-values correspond to the association or between-group comparison for each academic performance indicator. Test statistics other than correlation coefficients were removed to improve readability. \* $p < 0.05$ , \*\* $p < 0.01$ , \*\*\* $p < 0.001$ .

### 3.4. Sleep Quality and Academic Performance

Overall, sleep variables showed a non-uniform pattern of associations across academic indicators, with some results suggesting contradictory relationships depending on the outcome considered. Table 4 illustrates the associations between various sleep metrics and academic performance.

**Table 4.**
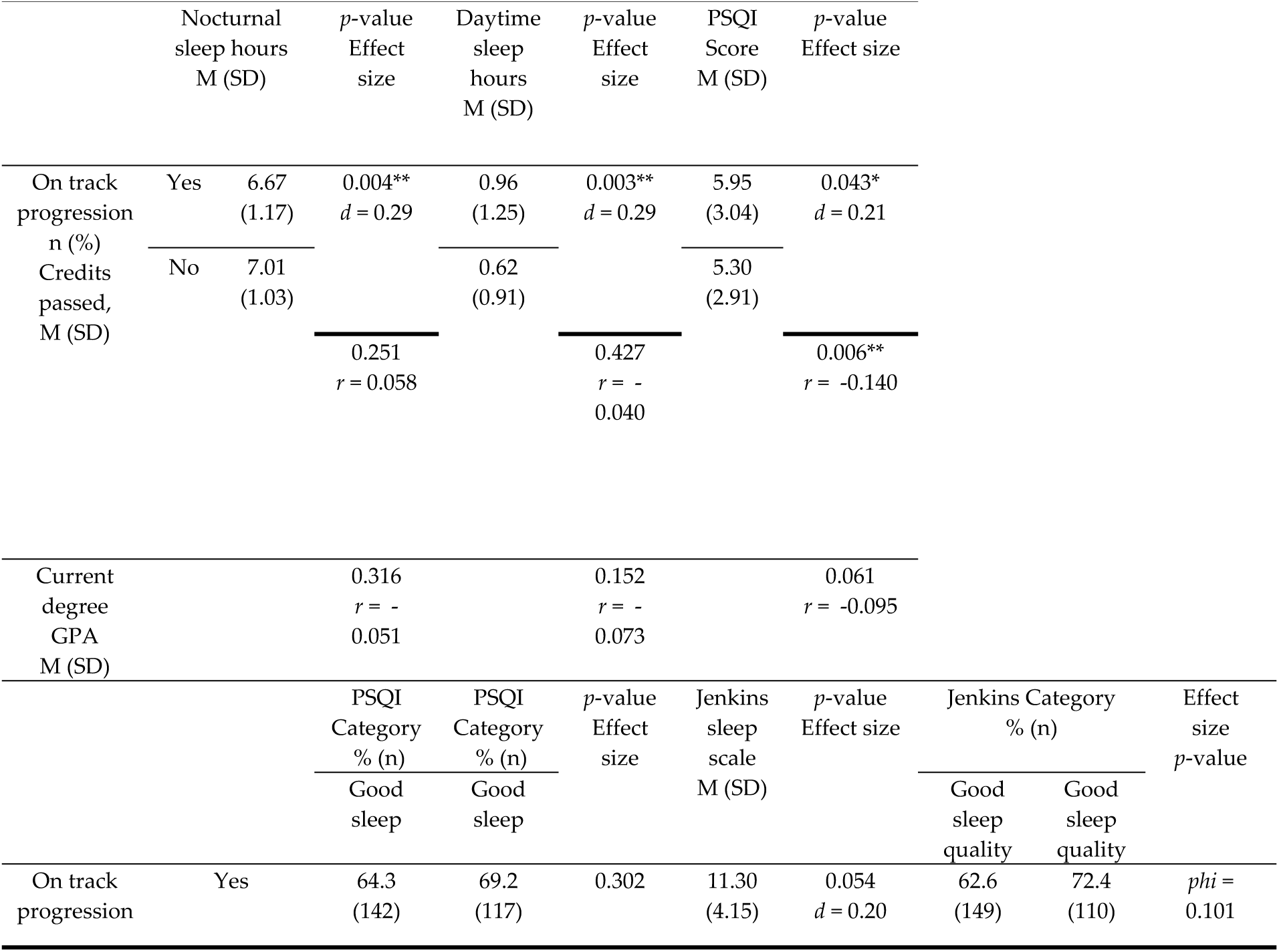
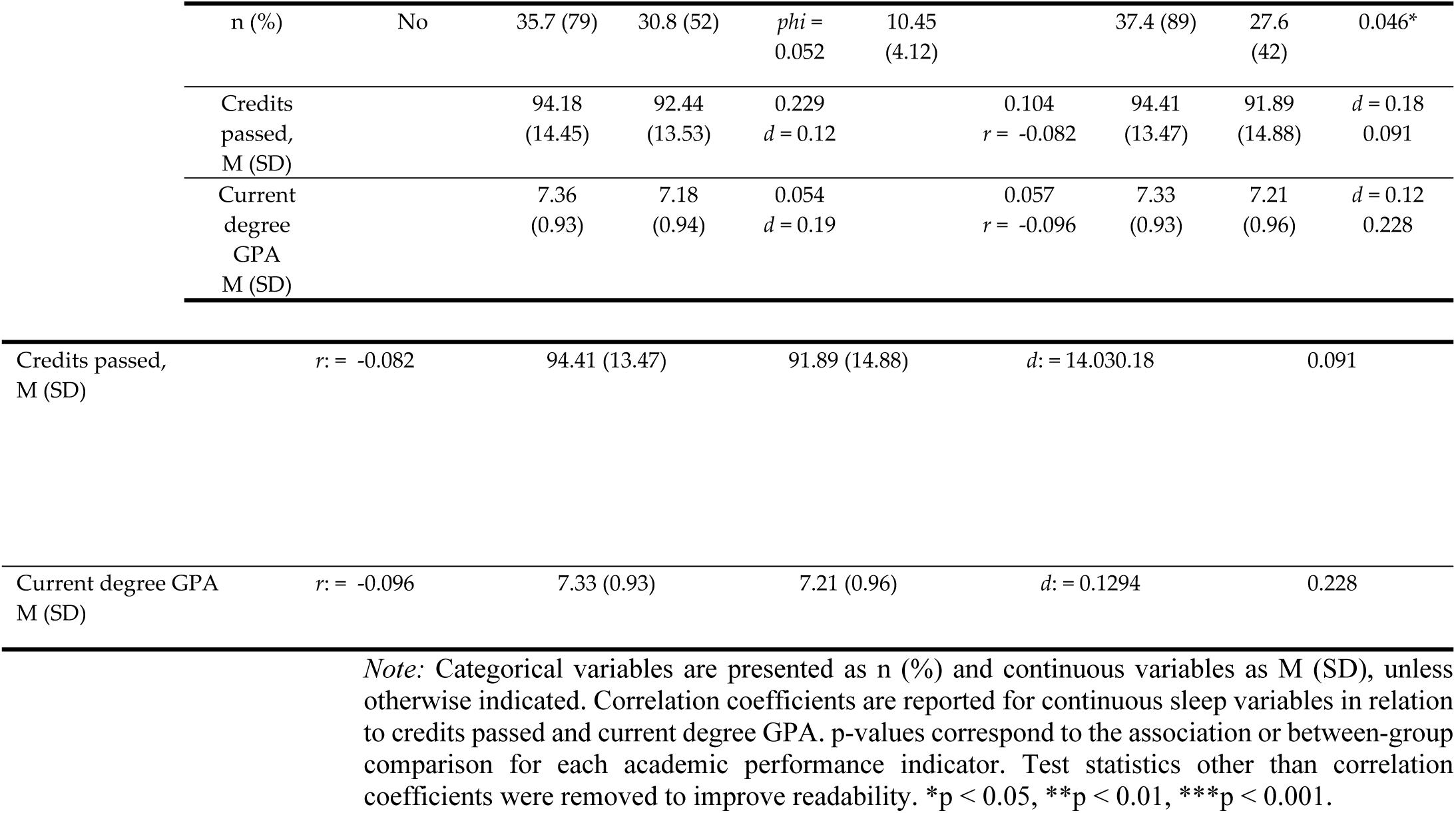
Relationship between sleep quality and academic performance.

Regarding academic progression, paradoxical results were observed. Students who advanced without repeating a year exhibited poorer sleep quality as measured by the PSQI (*M* = 5.95 vs 5.30; *t* = 2.02, *p* = .043) and the Jenkins Scale (*χ*² = 3.96, *p* = .046; ASR = ±2), with a small effect size (*phi* = 0.101). Furthermore, this group reported fewer hours of nocturnal sleep (*t* = 2.90, *p* = .004) but a higher number of daytime sleep hours (*t* = 3.03, *p* = .003), with a small effect size (*d* = 0.29), compared to those who did not progress at the expected rate.

In terms of continuous academic performance, the percentage of credits completed showed a significant negative correlation with PSQI scores (*r* = −.140, *p* = .006), indicating a small effect size *(d* = 0.12). Given that higher PSQI scores indicate greater sleep dysfunction, this result confirms that poor sleep quality is associated with lower success in credit attainment. No significant differences were found for grade point average (GPA), although marginal trends were observed for both the PSQI (*p* = .054) and Jenkins scale (*p* = .057). This suggests that better rest may be linked to higher grades, though these results did not reach the conventional threshold for statistical significance.

### 3.5. Physical Health, Smoking, and Academic Performance

Table 5 summarizes the relationships between smoking habits, physical health indicators, and academic performance.

**Table 5.**
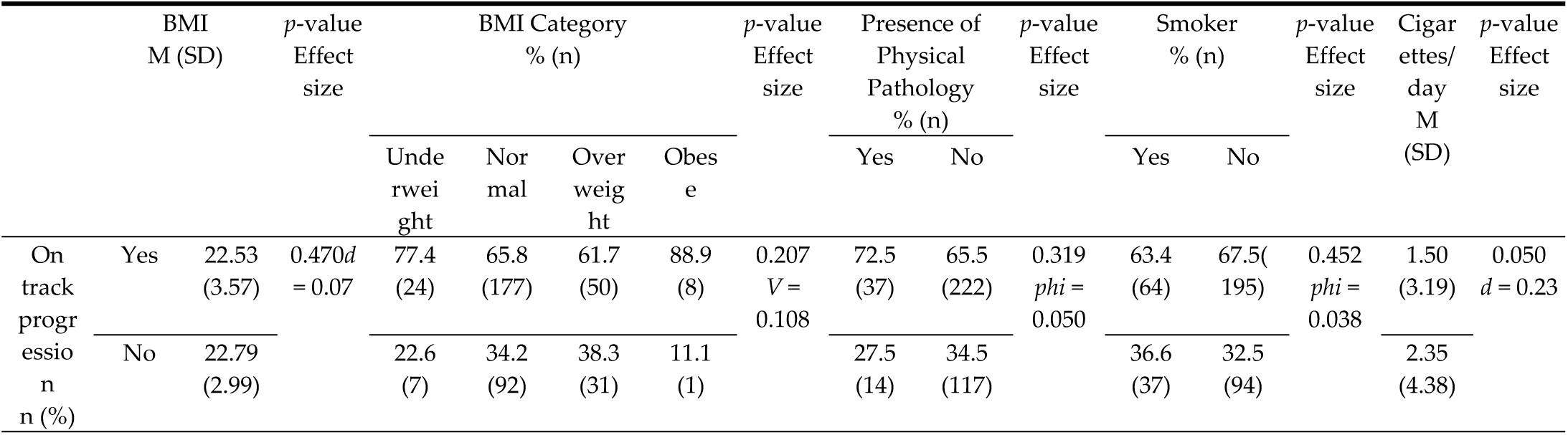
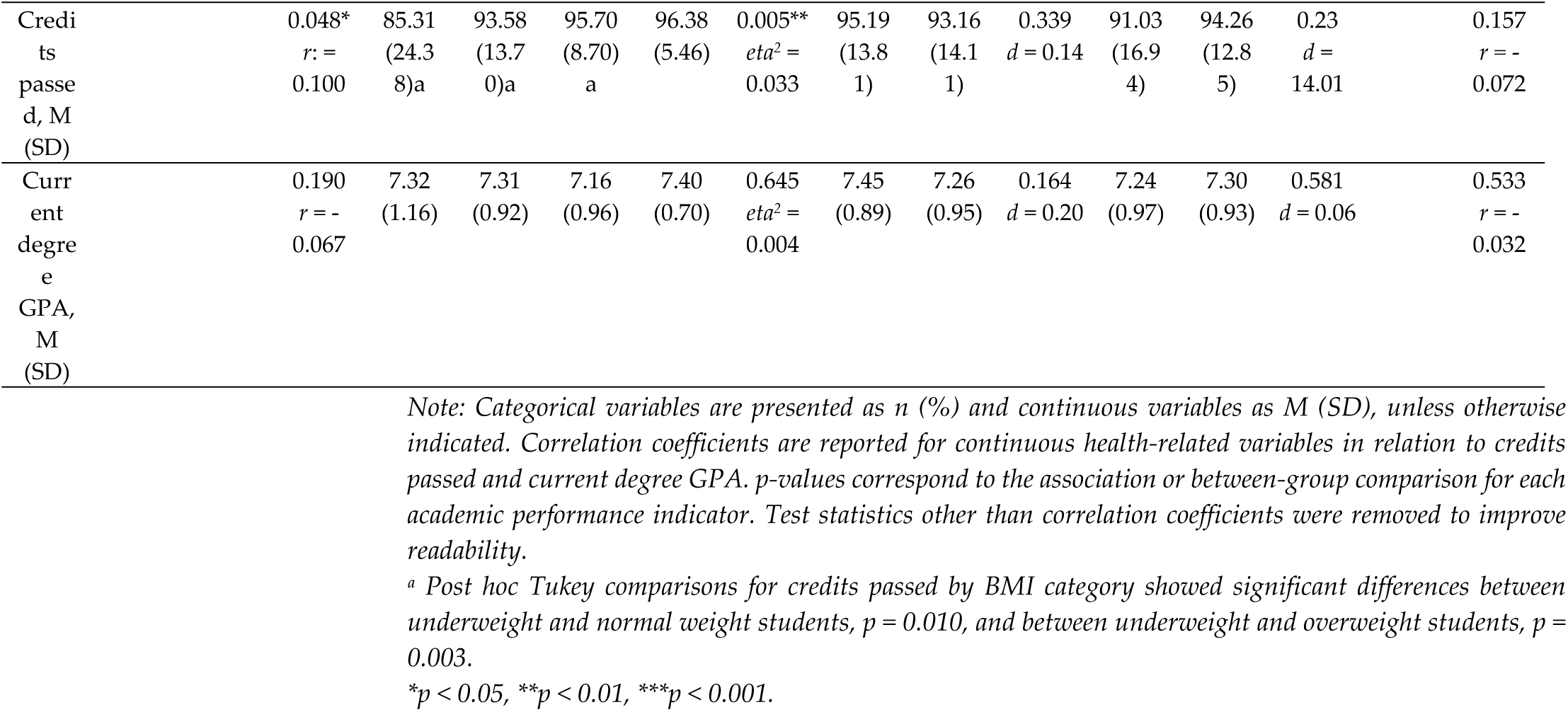
Relationship between smoking, physical health, and academic performance.

No significant associations were observed between tobacco consumption and academic performance (*p* > .05). However, a marginal difference was identified regarding the number of daily cigarettes, where students who did not progress through the academic year tended to smoke more than those who did (*t* = 1.96, *p* = .050), with a small effect size *(d* = 0.23).

Regarding BMI, a weak positive correlation was detected with the percentage of credits completed (*r* = .100, *p* = .048), with a small effect size, indicating that higher BMI values were associated with a higher rate of credits passed. Analysis of variance (*F* = 4.39, *p* = .005), with a small effect size (*eta*² = 0.033), and Tukey’s post-hoc tests indicated that underweight students completed a significantly lower percentage of credits compared to those with overweight (*p* = .010) or obesity (*p* = .003)

No significant associations were found between BMI and mean GPA. Similarly, the presence of physical pathology showed no significant relationship with any of the academic performance indicators analyzed.

### 3.6. Multivariate and Structural Modelling

To evaluate the predictive capacity of the variables that showed significant associations with academic performance, Generalised Linear Models (GLM) were employed. This analytical approach was selected as the most appropriate method given the heterogeneous nature of the study’s indicators, which included a mix of categorical and continuous variables. The use of GLMs provides greater statistical stability and robustness in parameter estimation for this specific data structure. The adjusted GLMs (Table 6) revealed several significant predictors. Figure 1 illustrates the structural relationships between lifestyle, health, and academic performance variables.

**Figure 1.**
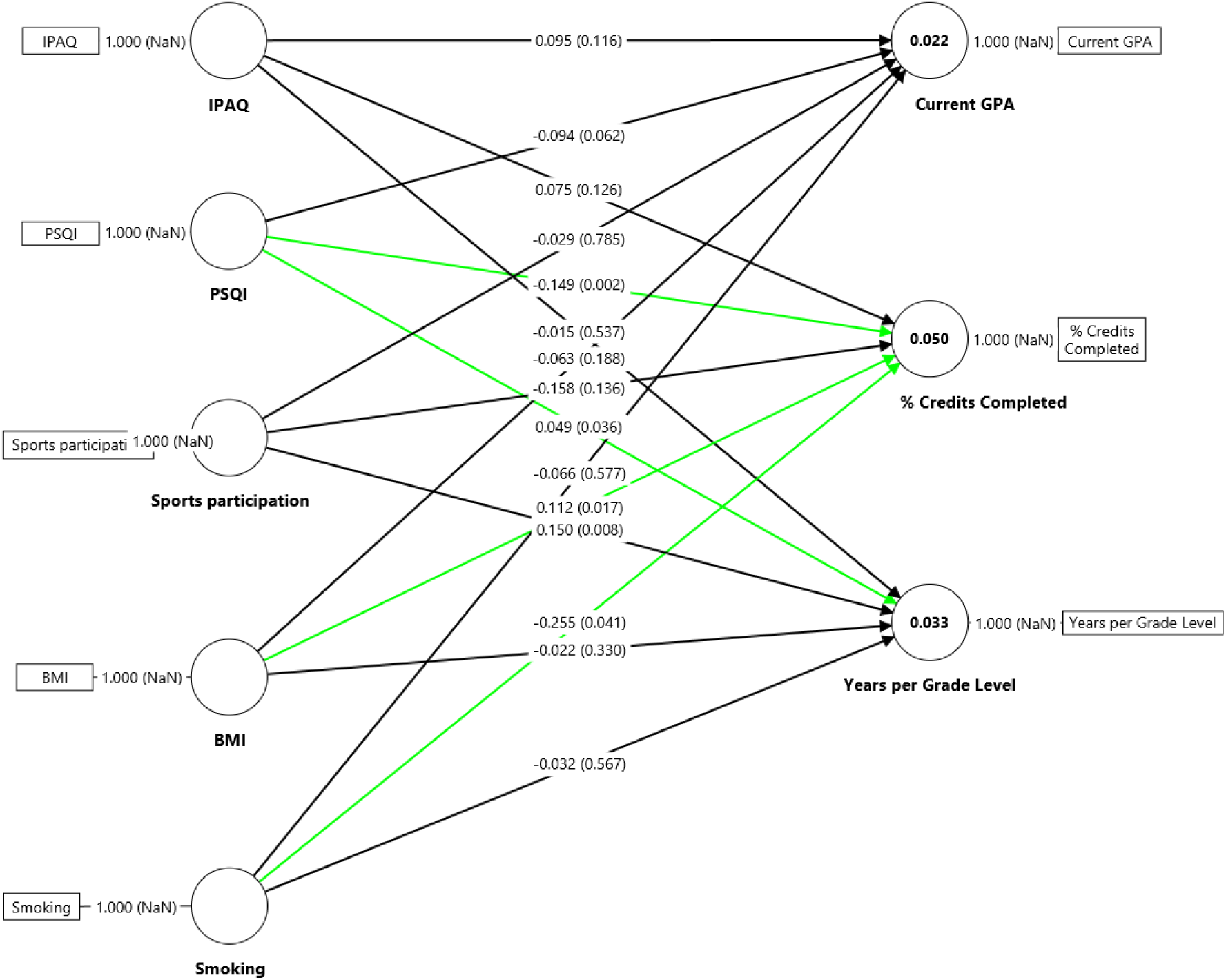
Structural model of the relationships between physical activity (IPAQ), sleep quality (PSQI), BMI, smoking, and academic performance indicators (GPA, percentage of credits completed, and years per grade level). **Note.** Standardised coefficients (β) are presented. Arrows indicate direct relationships between variables. Values in parentheses represent p-values. Green lines indicate statistically significant associations (p < .05). *R² = .022 for GPA, R² = .050 for percentage of credits completed, and R² = .033 for years per grade level.* IPAQ = International Physical Activity Questionnaire; PSQI = Pittsburgh Sleep Quality Index; BMI = Body Mass Index.

**Table 6.**
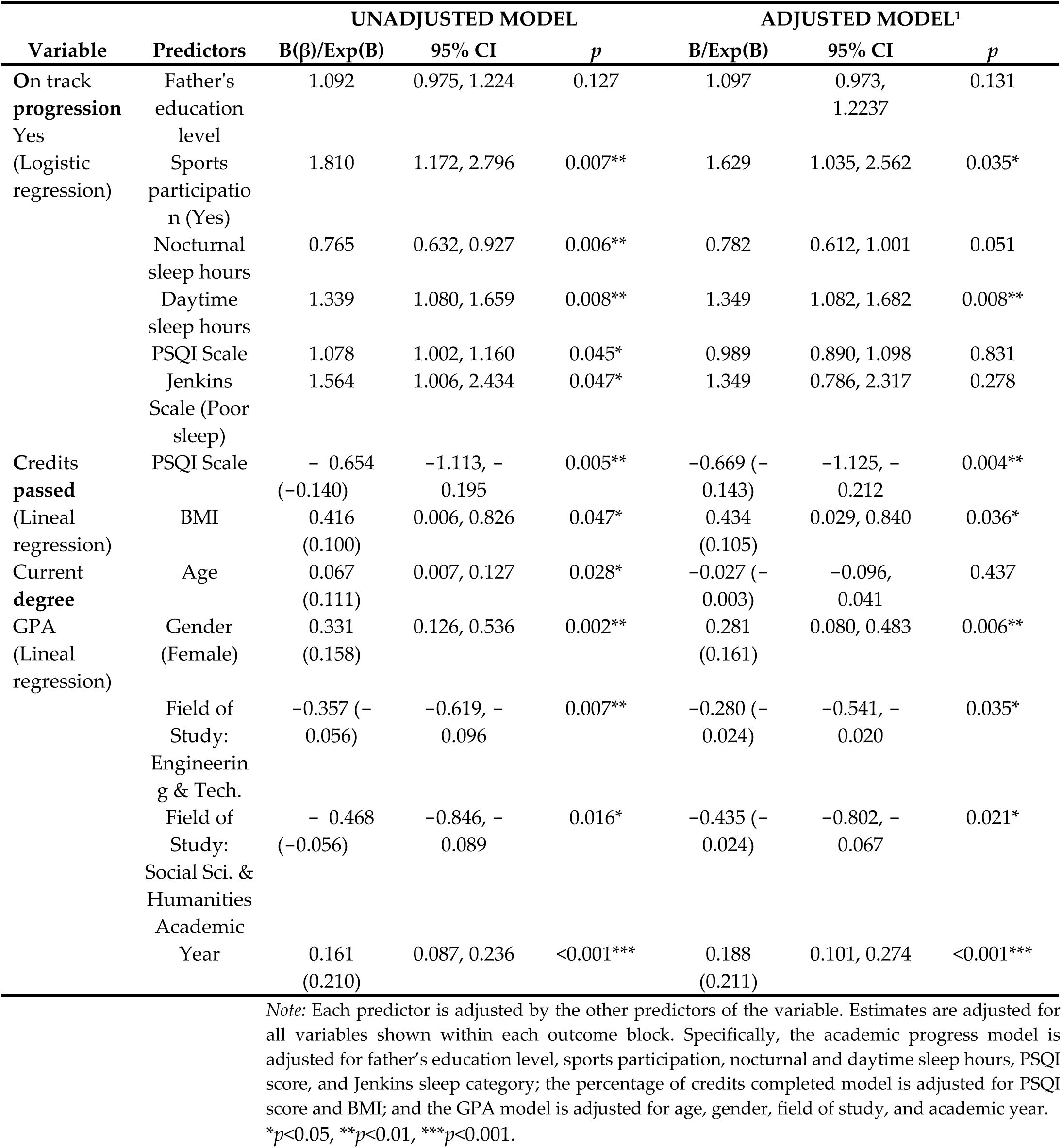
Generalised Linear Models for Predicting Academic Performance.

Students who participated in sports regularly were more likely to progress without repeating a year (Exp(B) = 1.629, *p* = 0.035), indicating a small-to-moderate effect in practical terms. Likewise, a higher number of daytime sleep hours significantly increased the probability of remaining on track with the academic curriculum (Exp(B) = 1.349, *p* = 0.008), reflecting a small effect size.

The percentage of credits completed was significantly explained by sleep quality, as measured by the PSQI (Exp(B) = 0.512, *p* = 0.004), indicating a moderate effect size, and by BMI (Exp(B) = 1.544, *p* = 0.036).

Overall, in this specific sample, poorer sleep quality and higher BMI were associated with a higher percentage of completed credits in this sample, indicating a small-to-moderate effect size; however, this finding should be interpreted cautiously and does not imply a beneficial effect of increased body weight.

Several factors predicted mean GPA. Female gender was associated with higher grades (Exp(B) = 1.325, *p* = 0.006), reflecting a small effect size. Regarding the field of study, students in Engineering and Technology (Exp(B) = 0.755, *p* = 0.035) and Social Sciences and Humanities (Exp(B) = 0.648, *p* = 0.021), both reflecting small-to-moderate effects, showed significant differences in GPA compared to other branches. Finally, the academic year was a significant predictor (Exp(B) = 1.206, *p* < 0.001), indicating a small-to-moderate effect size, suggesting that students in more advanced years tended to achieve higher grades.

As shown in Figure 1, most direct effects on current GPA were small and non-significant, with only weak contributions from the studied predictors (R² = 0.022). In contrast, several variables showed significant associations with intermediary academic outcomes. Sleep quality (PSQI) demonstrated a significant negative association with the percentage of credits completed (β = −0.149, p = 0.002), indicating that poorer sleep quality was linked to lower academic progress. BMI also showed a significant positive relationship with the percentage of credits completed (β = 0.112, p = 0.017), although the effect size was small.

Regarding academic progression (years per grade level), sports participation (β = 0.049, p = 0.036) and smoking (β = −0.255, p = 0.041) emerged as significant predictors, with sports participation associated with better progression and smoking associated with poorer progression. Additionally, BMI showed a small but significant positive effect on progression (β = 0.150, p = 0.008).

No significant direct effects were observed for IPAQ on any academic outcome, and most remaining paths were non-significant, confirming that the overall explanatory power of the model was limited (R² = 0.050 for percentage of credits completed and R² = 0.033 for years per grade level). Overall, the model suggests that lifestyle and health-related variables exert modest and selective effects on academic performance, primarily through academic progression and credit completion rather than GPA.

## 4. Discussion

The results of the present study partially confirm the proposed hypotheses and provide relevant evidence regarding the relationship between specific lifestyle habits and university academic performance. Collectively, it was observed that sleep habits and regular physical activity were significantly associated with academic progression, operationalised as advancing one year per grade, and completing a higher percentage of credits. However, while sleep demonstrated consistent associations with various indicators of academic success, physical activity was not significantly related to the percentage of credits completed. These patterns are further supported by the structural model (Figure 1), which shows that lifestyle variables exert modest and selective effects on academic outcomes, with low explained variance and primarily indirect pathways through academic progression and credit completion rather than GPA.

These findings suggest that physical exercise may primarily foster continuity and persistence within the curriculum, whereas sleep habits appear to have a more cross-cutting impact on academic performance. Conversely, GPA was more strongly associated with sociodemographic variables such as age, sex, and the specific degree program. This aligns with previous research indicating that personal and contextual factors differentially modulate quantitative indicators of academic achievement (Rodríguez-Hernández et al., 2026). The strong gender effect observed in GPA contrasts with the relatively small effect of field of study, suggesting that individual-level factors may play a more prominent role than academic context in explaining performance differences in this sample. No significant interaction effects were observed, reinforcing the consistency of these associations across sociodemographic groups. This interpretation is consistent with the structural model (Figure 1), which showed a very low explained variance for GPA (R² = .022), reinforcing the notion that it is influenced by factors beyond the lifestyle variables included in this study, and more strongly driven by sociodemographic characteristics.

The findings related to sleep partially align with existing literature, which highlights its importance for learning and cognitive functioning. Previous studies have shown that sleep quality is significantly associated with academic performance, while sleep duration yields less consistent results (Seoane et al., 2020). Furthermore, a recent meta-analysis demonstrated that sleep restriction negatively affects memory, showing a small but consistent effect (Hedges’ g = 0.29), which supports the plausibility of a cognitive impact of sleep on academic outcomes (Crowley et al., 2024).

However, the results of the present study reveal a mixed and partially contradictory pattern, as different sleep indicators were associated with academic performance in opposite directions depending on the outcome measure. Paradoxically, students with a higher probability of adequate academic progression exhibited poorer nocturnal sleep quality and shorter nighttime rest duration, accompanied by a higher number of daytime sleep hours. This pattern suggests that, within this sample, academic advancement may be associated with a redistribution rather than an absolute deficit of sleep, where reduced nocturnal sleep is partially compensated by daytime sleep. In line with this interpretation, previous research has shown that compensatory napping may mitigate some of the cognitive deficits associated with insufficient nocturnal sleep, particularly in young adults (Cousins et al., 2019; Leong & Chee, 2023; Milner & Cote, 2009). Nonetheless, this interpretation should be approached with caution, given the cross-sectional nature of the study.

In addition, this pattern may reflect the influence of chronotype and academic demands. University students frequently present an evening chronotype, which is associated with shorter nocturnal sleep due to early academic schedules and increased daytime sleepiness (Hershner & Chervin, 2014). Under these conditions, students may adopt compensatory strategies, such as daytime sleep or flexible study schedules, that allow them to maintain academic progression despite suboptimal sleep quality (Barley et al., 2023). At the same time, it is plausible that students who remain on track allocate more time to academic tasks at the expense of nocturnal sleep, particularly during high-demand periods. This suggests a potential "Academic Trade-off" between sleep and academic workload (Alblewi et al., 2025). It is crucial to clarify that these results do not suggest that sleep deprivation is beneficial for learning or cognitive function; rather, the increase in daytime sleep hours observed in this group should be interpreted as an academic survival strategy. Students appear to utilise napping as a compensatory mechanism to mitigate the neurocognitive deficits resulting from nocturnal sleep fragmentation.

In this regard, the observed association between poorer sleep quality and a higher percentage of completed credits contrasts with studies that have found robust relationships between superior sleep patterns and higher university grades, particularly when using objective measures such as wearable devices (Okano et al., 2019). A plausible explanation lies in the presence of compensatory effects, whereby students with poorer sleep quality increase the time dedicated to studying at the expense of rest. Alternatively, unmeasured variables such as academic stress, chronotype, or learning strategies may play a role. In fact, other studies have failed to find significant differences in academic performance based on sleep quality (Jalali et al., 2020), suggesting that this relationship is not uniform and may depend on individual and contextual factors. Importantly, these findings may also help explain the differential role of sleep quality across outcome variables in the multivariate models. While PSQI significantly predicted the percentage of credits completed, it was not a significant predictor of academic progression. This discrepancy may be due to the nature of the indicators: academic progression is a dichotomous and relatively coarse measure (on track vs. delayed), likely influenced by behavioral consistency and persistence, whereas the percentage of credits completed constitutes a continuous and more sensitive indicator of academic productivity, more directly affected by cognitive processes such as attention, memory, and fatigue, which are known to be influenced by sleep quality (Curcio et al., 2006; Leong & Chee, 2023). These findings suggest that the relationship between sleep and academic performance may differ depending on the specific academic indicator considered, reflecting distinct underlying mechanisms.

Regarding physical activity, the results indicate that regular practice is positively associated with academic progress, in alignment with previous evidence. Various studies have noted that physical exercise contributes to emotional regulation, attention, and executive functions, key processes for academic functioning (Zhao et al., 2024). Among university populations, longitudinal studies have shown that physical activity is associated with better stress recovery and variables linked to academic performance (Teuber et al., 2024), findings that have also been observed in secondary school students (Martín-Bozas et al., 2025). Consistently, a recent systematic review identified a moderate positive effect of exercise on academic performance in university students (Rosales-Ricardo & Cáceres-Manzano, 2024). These findings concur with research indicating that the impact of physical exercise is particularly robust regarding self-regulation, motivation, and executive functions (Zeng et al., 2017), thereby facilitating academic persistence and adaptation to the university environment. Furthermore, longitudinal studies have shown that regular physical activity predicts greater emotional stability and academic continuity, although it does not always translate into direct improvements in final grades (Kwan et al., 2012). This evidence underscores the notion that the benefits of physical exercise are more clearly reflected in indicators of academic consistency and progression than in strictly quantitative measures of performance. Acknowledging this distinction is vital for the field, as research has traditionally centered on GPA to measure academic performance. Rather than serving as an alternative, academic progression acts as a complementary metric that may help reconcile mixed results in literature. This perspective suggests that while exercise might not always lead to immediate grade increases, it supports the foundational consistency and persistence necessary for students to remain on their academic trajectory.

Regarding BMI, a modest association with the percentage of credits completed was observed. Previous evidence indicates that the relationship between BMI and academic performance is typically weak but consistent. He et al. (2019) analysed a sample of over 160,000 students and found a small negative correlation (r = -0.111) between these two variables. Nonetheless, this association tends to be attenuated when other health and lifestyle factors, such as nutrition, physical activity, or sleep, are controlled for. This suggests that BMI is better understood as an indirect marker than as a causal predictor of academic performance.

Within university populations, previous studies have identified negative correlations between BMI and final grades (Anderson & Good, 2017), as well as associations between BMI and lifestyle patterns that influence both general well-being and academic achievement. In contrast to much of this literature, BMI emerged as a small but significant positive predictor of the percentage of credits completed within the present sample. This finding must be interpreted with caution. It does not imply that a higher BMI is protective or beneficial for academic performance. Categorical analysis indicated that this positive association is largely attributable to the underweight group, whose performance was significantly lower than that of students in the overweight and obese ranges. Although these factors were not assessed in the present study, being underweight could reflect lower energy reserves or nutritional insufficiencies that might hinder the ability to cope with academic demands.

These findings could suggest that universities might benefit from broadening their approach to promoting healthy habits. Recent research (Gunsalus et al., 2024) warns that traditional medical nutritional guidelines often omit key determinants, which can exacerbate weight stigma and increase the risk of eating disorders. While previous literature has focused predominantly on obesity, the present results suggest that, at least in this sample, the underweight group accounted for the observed effect. Future research should determine whether this pattern is replicable in comparable samples. If confirmed, university health programs may benefit from broadening their scope to address underweight and not only overweight.

Another relevant finding was that no significant associations were observed between tobacco consumption and academic variables. Although the overall sample size was sufficient to detect small-to-moderate effects, the statistical power to identify associations in specific subgroups, particularly for variables with lower variability such as tobacco consumption, may have been limited. Therefore, non-significant findings in these cases should be interpreted with caution. This lower prevalence might have also been due to the sample being predominantly composed of women, which previous studies in the same Spanish population have already shown smoke less than men (Carmona Simarro et al., 2019). Nevertheless, previous studies have linked smoking to attentional difficulties and reduced concentration (Teixeira-da-Costa et al., 2022), as well as lower academic performance and higher absenteeism (Alqahtani et al., 2023). Furthermore, other studies employing similar single-item measures have yielded significant results; for instance, Mahfouz et al. (2024) found that students who smoked six to ten cigarettes per day were more likely to have a lower GPA, a correlation not observed in our findings. Moreover,, research in adolescent populations has found that the use of both conventional and electronic cigarettes is associated with lower academic achievement (Sheikhattari et al., 2025). The shift in consumption toward alternative nicotine delivery systems, such as e-cigarettes, may reduce the statistical power to detect associations based solely on combustible tobacco. Recent studies have highlighted that the prevalence of dual and exclusive e-cigarette users is increasing among European students (World Health Organisation, 2025) and that these modalities may present distinct cognitive profiles compared to traditional smokers (Teixeira-da-Costa et al., 2022).

Taken together, the results indicate that sleep habits, physical activity, and, to a lesser extent, BMI, constitute relevant variables in explaining academic progress, whereas tobacco consumption appears to hold less weight in this population. These findings reinforce the need to promote university health policies and programs aimed at improving sleep, physical activity, and nutrition, incorporating an approach that addresses both underweight and overweight prevention.

Overall, the structural model reinforces the interpretation that lifestyle and health-related variables play a secondary but meaningful role in academic performance, with their influence being more evident in behavioural indicators of academic continuity than in final grades, and largely operating through indirect pathways.

### Limitations and further research

The study presents some limitations. The cross-sectional design precludes causal inferences regarding the direction of the observed associations. The use of self-reports may have introduced recall or social desirability biases. In addition, the sampling procedure combined two recruitment phases: an in-person street-level approach in public areas near universities in Valencia and a subsequent online distribution of the same questionnaire. Although the same eligibility criteria and questionnaire were applied in both phases, the combination of in-person and online recruitment strategies may have introduced heterogeneity and potential selection bias, as participants recruited through different modalities may differ in unobserved characteristics. A relevant limitation concerns the gender imbalance in the sample, with women representing 71.3% of participants. This disproportion may have influenced the results, particularly those related to academic performance, as female students showed significantly higher GPA scores with a small-to-moderate effect size. Consequently, the overall estimates may be partially driven by this overrepresentation, potentially inflating associations linked to variables correlated with gender. Although gender was included as a covariate in the multivariate models, this statistical adjustment does not fully eliminate the potential bias derived from unequal group sizes. Therefore, the findings should be interpreted with caution, especially when generaliszing to more gender-balanced populations. Future studies should aim to achieve more balanced sampling to better disentangle gender-specific effects and improve external validity. Regarding the IPAQ, the average score was higher than typically reported in university student populations. Notably, 12 participants (approximately 3% of the sample) reported values above 10,000 MET-min/week. Because these observations represent a meaningful proportion of the sample and correspond to individuals with particularly high levels of physical activity, they were retained in the analyses. Nevertheless, their presence may have contributed to the elevated mean score and should be considered when interpreting the findings. Additionally, the study omitted different variables that could be confounding the results and are of interest in relation to academic achievement. Our sociodemographic characteristics lacked relevant socioeconomic data that is also associated with academic performance such as prior academic achievement, university experience and working status (Rodriguez-Hernandez et al., 2020), which could be confounding our results. Furthermore, relevant psychosocial variablesuch as perceived stress, anxiety, social support, and study habits as well as potential differences in curricular workload, were not included; these factors could act as mediators or moderators of the associations between lifestyle and academic performance (Richardson et al., 2012). Lastly, alcohol consumption was not included in the present study due to the need to limit questionnaire length and reduce participant burden; however, given its established association with academic performance, it should be considered in future research. Additionally, tobacco consumption was assessed using two ad hoc items, which, although practical for brief screening, do not capture other relevant patterns such as vaping, dual use, frequency or intensity of consumption, or nicotine dependence, thereby limiting the interpretability of these findings. Finally, although BMI was included as an accessible anthropometric indicator, in young adults, it may not accurately reflect body composition, since individuals with similar BMI values may differ in fat mass, lean mass, and fat distribution. This may have limited the precision of interpretations involving weight status in the present study.

Nonetheless, the results of the present study have significant implications for the university setting. The promotion of sleep hygiene programs, the encouragement of regular physical activity and health education, and the reduction of harmful habits could contribute significantly to improving both academic performance and student mental health. Universities could integrate these actions into counselling and tutoring services or educational campaigns, fostering healthier academic environments and the development of self-care strategies.

This study suggests that university health policies and programs aimed at improving nutrition should also address underweight prevention as well as overweight prevention and a healthy diet. Additionally, our findings also underscore the notion that the benefits of physical exercise are more clearly reflected in indicators of academic consistency and progression than in strictly quantitative measures of performance. These findings might also aide to interpret some of the inconsistencies in the literature regarding the association between physical activity and academic performance. Furthermore, our evidence regarding nicotine consumption in Spanish university women supports Carmona Simarro et al.’s (2019) evidence. The lack of significant results in our study also highlights the importance of future research on tobacco to pay special consideration to sampling both genders equally. Additionally, in some lifestyle habits interventions, such as improving dietary intake, gender-targeted interventions have been shown to be more effective than gender-neutral interventions (Sharkey et al., 2020). Thus, perhaps university programs promoting healthy lifestyles related to nicotine consumption could benefit from targeting this specific program for men to achieve better results. Moreover, this study also highlights the importance of paying attention to alternative nicotine devices.

For future research, it is recommended to employ longitudinal or experimental designs and incorporate objective measures of sleep and physical activity (e.g., via actigraphy and accelerometry). Recent studies have utilised actigraphy to explore multiple dimensions of sleep and their association with academic outcomes, highlighting that sleep variability and onset timing may be associated with poorer academic performance, even when controlling for other relevant variables (Mathew et al., 2024). Additionally, future research could benefit from using different measures of academic performance, such as academic progression, as it might help further understand the associations between health habits and academic performance, especially in the case of physical activity. Furthermore, it is suggested to explore mediating and moderating variables such as self-regulated learning, academic motivation, and fatigue. They have been shown to play a key role in the relationship between psychological processes and academic outcomes, even acting as intermediate variables in the link between motivation and academic procrastination (Aulia et al., 2024). Likewise, given the rising prevalence of alternative nicotine consumption, such as vaping, among youth and its potential impact on attention, behaviour, and academic outcomes, it is pertinent to specifically examine the impact of vaping on academic performance and study habits, particularly as epidemiological studies have found associations between e-cigarette use and poorer school outcomes (Augenstein et al., 2024). Additionally, when researching the effects of nicotine consumption, studies should pay special consideration to sampling both genders equally. As perhaps university programs promoting healthy lifestyles related to nicotine consumption could benefit from targeting this specific program for men. Finally, future work could examine potential differences by sex, field of study, and other contextual factors (e.g., curricular demands or technology use patterns) that may influence the mechanisms connecting lifestyle to academic performance.

## 5. Conclusions

This study demonstrates that specific lifestyle habits are differentially associated with academic performance in university students, with clear directionality depending on the outcome considered.

First, sleep patterns showed consistent and significant associations with academic performance, although the direction of these associations varied by indicator. Notably, poorer nocturnal sleep quality and shorter nighttime duration were associated with better academic progression and higher percentages of completed credits, a pattern likely reflecting compensatory strategies such as increased daytime sleep. This finding suggests that, in this population, academic success is linked to a redistribution of sleep rather than optimal sleep quality, highlighting a trade-off between academic demands and nocturnal rest.

Second, regular physical activity was positively associated with academic progression, indicating that students who engage in exercise are more likely to remain on track in their studies. However, physical activity was not associated with GPA, suggesting that its benefits are more strongly related to persistence and continuity than to quantitative academic achievement.

Third, BMI showed a small but significant positive association with the percentage of credits completed, primarily driven by lower performance among underweight students, rather than a beneficial effect of higher BMI. These finding positions underweight status as a potential risk marker for poorer academic outcomes.

Fourth, sociodemographic factors (particularly gender and academic year) were stronger predictors of GPA than lifestyle variables, indicating that quantitative performance is more influenced by individual and contextual characteristics than by health behaviours.

Overall, these findings indicate that lifestyle factors play a modest but meaningful role in academic performance, primarily influencing behavioural indicators such as progression and persistence rather than grades, and often through indirect or compensatory mechanisms.

## Author Contributions

Conceptualization, P.S. and P.P.; methodology, T.L.; software, A.B.; validation, P.S., A.B. and T.L.; formal analysis, A.B.; investigation, P.S.; resources, B.T.; data curation, A.B.; writing—original draft preparation, P.S and P.P.; writing—review and editing, B.T.; P.P.; visualization, B.T.; supervision, G.H; P.P.; project administration, G.H. All authors have read and agreed to the published version of the manuscript.

## Funding

This research received no external funding.

## Institutional Review Board Statement

The study was conducted in accordance with the Declaration of Helsinki, and approved by the Ethics and Biomedical Research Committee of Universidad Cardenal Herrera-CEU (Ref: CEEI23/440, date of approval 19 February 2024).

## Informed Consent Statement

Informed consent was obtained from all subjects involved in the study.

## Data Availability Statement

The data presented in this study are available from the corresponding author upon reasonable request. The data are not publicly available due to privacy and ethical restrictions.

## Acknowledgments

The Group of Investigation TXP was fundamental to developing this project.

## Conflicts of Interest

The authors declare no conflicts of interest.

## Abbreviations

The following abbreviations are used in this manuscript:

ANOVA: Analysis of variance
ASR: Adjusted standardized residuals
BMI: Body Mass Index
CI: Confidence interval
GLM: Generalized Linear Models
GPA: Grade Point Average
HL: Healthy Lifestyle
IPAQ: International Physical Activity Questionnaire
JSS-4: Jenkins Sleep Scale-4
M: Mean
METs: Metabolic Equivalent of Task
PSQI: Pittsburgh Sleep Quality Index
SD: Standard Deviation
WHO: World Health Organization

## Notes

### Competing Interest Statement

The authors have declared no competing interest.

